# Compact and evenly distributed *k*-mer binning for genomic sequences

**DOI:** 10.1101/2020.10.12.335364

**Authors:** Johan Nyström-Persson, Gabriel Keeble-Gagnère, Niamat Zawad

## Abstract

The processing of *k*-mers (subsequences of length *k*) is at the foundation of many sequence processing algorithms in bioinformatics, including *k*-mer counting for genome size estimation, genome assembly, and taxonomic classification for metagenomics. Minimizers - ordered *m*-mers where *m < k* - are often used to group *k*-mers into bins as a first step in such processing. However, minimizers are known to generate bins of very different sizes, which can pose challenges for distributed and parallel processing, as well as generally increase memory requirements. Furthermore, although various minimizer orderings have been proposed, their practical value for improving tool efficiency has not yet been fully explored. Here we present Discount, a distributed *k*-mer counting tool based on Apache Spark, which we use to investigate the behaviour of various minimizer orderings in practice when applied to metagenomics data. Using this tool, we then introduce the universal frequency ordering, a new combination of frequency counted minimizers and universal *k*-mer hitting sets, which yields both evenly distributed binning and small bin sizes. We show that this ordering allows Discount to perform distributed *k*-mer counting on a large dataset in as little as 1/8 of the memory of comparable approaches, making it the most efficient out-of-core distributed *k*-mer counting method available.

## Introduction

The analysis of *k*-mers, short sequence fragments of a fixed length, is foundational to many methods and algorithms in bioinformatics, including genome assembly [13], *k*-mer counting [7, 12, 20], variant calling [1], and metagenomic classification [23]. Due to the proliferation of next generation sequencing (NGS) data and other types of ‘omics data, such *k*-mer data analysis needs are constantly increasing. This has led to the need for ever more efficient algorithms and methods in this area. The *k*-mer analysis of large datasets is often computationally challenging. For example, when *k* = 55, for the usual DNA alphabet A, C, G, T there exists a total of 4^55^ (approx. 1.3 10^33^) possible such *k*-mers. This large data space, usually much too large to represent in memory or on disk in its entirety, is a major source of the complexity of *k*-mer analysis. One commonly used strategy for overcoming this complexity is *k*-mer binning. Since only a small fraction of all possible *k*-mers are seen in practice for a given dataset, one aims to subdivide the data into reasonably sized parts and base data processing (such as counting, manipulation, lookup of associated data) on these.

Binning is often done by dividing *k*-mers into *superkmers* according to their *minimizers*, a technique first introduced in biological applications by Roberts et al [21]. Minimizers are obtained by ordering all *m*-mers *M_i_* for some fixed *m*, where *m < k*, in some way: *M*_0_ *< M*_1_ *< … < M_n_*. We say that *M_i_* is smaller than *M_j_* if *i < j*. Each *k*-mer is then classified according to the smallest minimizer in it. Often, consecutive *k*-mers in a longer sequence will share the same *m*-minimizer. Thus, an input sequence may be split into superkmers, maximally long substrings where all *k*-mers share the same minimizer. In addition to partitioning the data into bins, where each bin corresponds to a minimizer, grouping *k*-mers together in this way can also provide a compact representation, since superkmers are much more space-efficient than representing each *k*-mer by itself.

Various minimizer orderings exist. Although simple to implement and reason about, the basic lexicographic ordering proposed by Roberts leads to very unbalanced bins in practice. This can lead to higher memory usage and to a slowdown in general, since algorithms on larger bins can be more expensive to run.

The *k*-mer counter KMC2 [4] introduced *minimizer signatures*, which order *m*-mers lexicographically, except that in order to reduce data skew, *m*-mers starting with AAA or ACA are given lower priority, and *m*-mers containing AA anywhere are also avoided, except for AA at the start. This helps spread out the *k*-mers somewhat and avoid certain very unbalanced bins. This ordering is also used by Gerbil [7], and by FastKmer [8] in a modified form with some additional rules.

The *frequency counted* ordering was first introduced by Chikhi et al [2] for the purpose of efficient de Bruijn Graph representation. A similar approach (weighted minimizers) was also used by Jain et al [10] for long read mapping. In this ordering, rare minimizers, based on occurrence in the actual dataset in each case, are given higher priority than common minimizers.

Finally, the concept of *compact universal k-mer sets* was recently introduced by Orenstein et al [17, 18]. For any given sequence of length *k* to be hit by (include) at least one sequence of length m in some set of *m*-mers, it is not necessary to include every *m*-mer in the set. Small sets that hit every *k*-length sequence can be precomputed. While computing minimal such sets is an NP-hard problem, heuristics can be used in practice to produce relatively small sets. Such a set can be turned into an ordering by giving all *m*-mers not in the set lower priority, effectively excluding them.

Although many different orderings with diverse characteristics have been proposed, it is still not clear which ordering should be preferred in practice, and many recent innovations have not yet been fully explored. Thus, to help identify the best methods for large scale ‘omics data analysis, a practical evaluation of the various possible orderings when applied to demanding tasks is needed.

To evaluate minimizer orderings, with a view to reducing memory usage and overall processing time, the following metrics are usually considered.

### Flatness of distribution

This can be measured in various ways, for example through the ratio of the maximum bin size to the mean bin size.

### Maximum bin size

Depending on the application, this may be even more important than flatness of distribution. For tools that perform sequential processing of bins, often the minimum memory requirement is that each bin should be able to fit in memory in its entirety.

### Length of superkmers/compactness

Longer superkmers give a more compact representation in memory and on disk, or for network transmission in a distributed setting. Equivalently, one may measure the average distance between minimizers (often called the density of minimizers), or the total number of superkmers (inversely proportional to their average length for a given dataset). ^1^

### Number of bins

Many tools currently in use also try to limit the number of bins. For example, the KMC2 authors argue that one goal of a good minimizer ordering should be that “the number of bins should be neither too large nor too small” [4]. However, while limiting the number of bins is reasonable when each bin is stored in a separate file, modern file formats allow a large number of bins to be stored together in a small number of files. Furthermore, a larger number of fine grained bins has advantages in subsequent processing. Many algorithms, such as sorting, generally have a lower per-item cost when applied to smaller lists of items, e.g. quicksort has a best case runtime of (*n* log *n*) and a worst case of (*n*^2^). Thus, in addition to the goals stated above, we believe that the ability to generate a large number of small (but evenly distributed) bins is a desirable goal, especially if this can also be done while keeping superkmers long.

Marçais et al provided a systematic study of the minimizer behaviour in practice of various widely used tools [15]. They compare existing tools with universal sets (DOCKS) for the parameters *m* = 7, *k* = 11, for a synthetic dataset as well as for the human genome. They reported the average bin size, the max/mean ratio for bins, and the mean distance between minimizers. The number of bins studied was in each case *<*= 16,384.

Erbert et al evaluated various minimizer orderings as part of their work on Gerbil [7], in the context of the F. vesca genome for *m* = 6, *k* = 28. In addition to the KMC2/signature ordering, lexicographic ordering, randomized ordering, and the CGAT (lexicographic, but with C *<* G *<* A *<* T) ordering, they also studied *distance from pivot* (dfp), which is a modified version of the frequency counted ordering that attempts to avoid very small bins. For this comparison, they only reported the maximum bin size and the total number of superkmers. Although compactness was measured, measurements such as the bin size distribution, or the number of bins generated in each case, were not reported.

In ‘omics data analysis, metagenomics data has much higher complexity compared to single-species ‘omics data, owing to the large number of distinct *k*-mers, and hence represents a challenging use case. Here we systematically study, using a metagenomics dataset, the use of frequency counted and universal set based minimizer orderings to generate a large number of evenly distributed *k*-mer bins. We consider cases in the order of 10^5^ to 10^6^ bins. As far as we know, this is the first time that these minimizer orderings have been comparatively evaluated for a large number of bins, or with metagenomics data. We then combine the two to construct a novel ordering, *frequency counted universal minimizers*, which yields a very even distribution and long superkmers.

With the increasing size and complexity of ‘omics datasets, and the high resource requirements of some ‘omics tools, there is an increasing need for distributed algorithms, which can run in a cluster or in the cloud. For distributed processing, being able to subdivide the workload evenly is even more important than for single-machine processing, since communication and synchronisation costs, such as network shuffling, can be significant. One of the more popular and widely accepted frameworks for distributed data processing in recent years is Apache Spark [22] (Spark for short), which is based on technologies such as Hadoop, but allows for a more general programming model than, for example, MapReduce. For Spark applications to be performant, being able to partition data evenly enough is very valuable [11].

As a case study for the performance benefits of minimizer orderings, we are particularly interested in the problem of *k*-mer counting [14]. In itself, this method can be used for purposes such as abundance filtering and genome size estimation, and it can also be a necessary foundation for more complex methods, such as de Bruijn Graph compaction, genome assembly, and metagenomic classification. Thus, in order to evaluate the various minimizer orderings, we implement a new distributed *k*-mer counting tool on Spark, called Discount. Discount can function as a pure *k*-mer counter, but can also double as a minimizer analysis tool, reporting detailed statistics about superkmers and *k*-mers in each bin, as well as supporting methods such as frequency sampling and user-supplied universal sets. This allows us to freely evaluate various orderings on a realistic workload. Discount is freely available (on GitHub at https://github.com/jtnystrom/discount) and GPL licensed.

## Methods

In order to study minimizer orderings as well as their effect on practical tasks, we have implemented a distributed *k*-mer counting tool, Discount, on Apache Spark. Here we briefly describe the design of this tool. Spark applications operate on *RDDs* (resilient distributed datasets), which are distributed collections of data, divided into some number of partitions. An application can execute some number of *stages* that operate on *partitions* of such RDDs in parallel on a cluster.

Figure 1 shows the internal stages of Discount. In order to read FASTA and FASTQ files efficiently into Spark, we use the FastDoop library [19]. Next, the following stages are applied.

**Figure 1.**
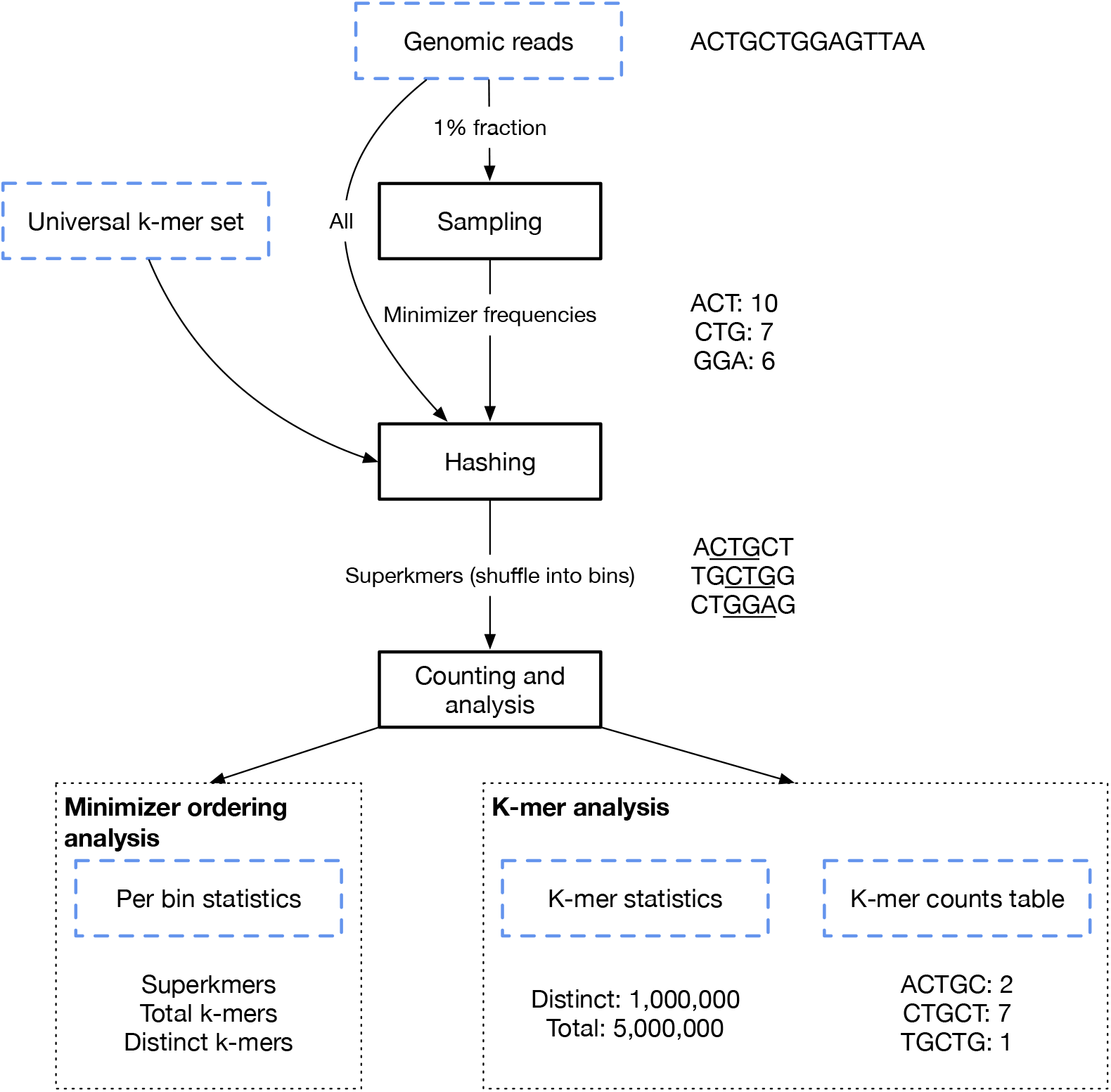
Internal stages of the Discount application. Here we show a toy example with *m* = 3 (minimizer length) and *k* = 5 (*k*-mer length)

### Sampling stage (optional)

When sampling is used, a fraction (our default is 1%) of reads is sampled to obtain an estimate of minimizer frequencies in the data.

### Hashing stage

Genomic reads are split into superkmers. If desired, the minimizer frequency estimate from the previous stage, and optionally also a user-supplied universal compact *k*-mer set, are used to configure the minimizer ordering. The resulting superkmers are encoded in the commonly used compact form of two bits per base pair.

### Counting and analysis stage

Superkmers are shuffled into their assigned bins (i.e., by minimizer) such that each bin is located in its entirety on one partition, and thus on a single machine in a Spark cluster.

Next, in *k*-mer counting mode, in each bin superkmers are broken up into individual *k*-mers, sorted, and counted. If users have requested a counts table or histogram to be output, then that is written to disk. Otherwise, only summary statistics is collected and aggregated, and then presented to the user. In minimizer analysis mode, a summary of the contents of bins is output in a table containing each bin’s number of superkmers, distinct *k*-mers, and total *k*-mers. This allows for further downstream evaluation of the selected minimizer ordering’s behaviour and characteristics.

We use Discount to study the following minimizer orderings.

### Minimizer signatures

We implement minimizer signatures according to the rules described in [4]. See the introduction for details.

### Frequency counted (sampled)

Here we order minimizers from rare to common based on their estimated abundance in the actual dataset. For efficiency, we sample 1% of the data and use frequencies obtained from this fraction. Ties between equally frequent minimizers are resolved by ordering them lexicographically.

### Universal sets

We use the precomputed DOCKS sets from [17], freely available at their website [16]. For this ordering, we exclude those minimizers that are not in the universal set, and the included minimizers are ordered lexicographically (A *<* C *<* G *<* T). We selected four sets to use in this work, shown in Table 1. For example, res_10_20_4_0 is a set of 10-mers guaranteed to hit every sequence of length 20 (or longer). This sets an upper limit to the number of bins that can be created in each case. We did not have access to precomputed sets for *k* = 28 and 55 specifically, and could not compute them ourselves due to the need for commercial solver software, but such sets would likely be smaller and yield more efficiency.

**Table 1.** Sizes of the minimizer sets used for the universal and universal frequency orderings in our evaluation.

| k | m | Set | m-mers |
| --- | --- | --- | --- |
| 28 | 10 | `res_10_20_4_0` | 214239 |
| 28 | 9 | `res_9_20_4_0` | 51591 |
| 55 | 10 | `res_10_50_4_0` | 128137 |
| 55 | 9 | `res_9_50_4_0` | 34458 |

### Universal frequency counted

We propose a novel ordering, obtained by combining the frequency counted ordering and the universal set ordering. This ordering is established by sorting DOCKS universal sets according to the 1% sampled frequency count in the data. As above, ties between equally frequent minimizers are resolved lexicographically.

## Results

The dataset being studied is a cow rumen metagenomic study [9], short read NGS data, available from the Sequencing Read Archive (SRA) as run SRR094926 from project PRJNA60251. In this section we use the first 100 million reads (Table 2). As a FASTQ file, this is approximately 30 GB.

**Table 2.** Size of the dataset used to evaluate minimizer orderings.

| k | Total $k$-mers | Distinct $k$-mers |
| --- | --- | --- |
| 28 | $7.40 \times 10^9$ | $5.09 \times 10^9$ |
| 55 | $4.70 \times 10^9$ | $3.73 \times 10^9$ |

Our collected bin statistics for the various orderings are shown in Table 3. The signature ordering has a high max/mean ratio, but produces long superkmers. The universal set ordering produces even longer superkmers for *k* = 28, but does not improve the max/mean ratio substantially. The frequency ordering greatly improves the max/mean ratio, but superkmers are much shorter. Finally, the universal frequency ordering produces the best max/mean ratio, and also produces almost as long superkmers as the signature ordering.

**Table 3.** Minimizer ordering measurements. Results obtained when using Discount to break up the dataset into binned superkmers. Here, *m* represents the minimizer length, so that for an unconstrained ordering, there would be a theoretical maximum of 4^*m*^ different minimizers (bins). Bin sizes are measured as the total number of *k*-mers in each bin prior to counting distinct items (i.e. as the sum of superkmer lengths in that bin). The average superkmer length is measured as a number of *k*-mers.

| k | m | Ordering | Bins | Bin size (total $k$-mers) | | | | Avg s.kmer |
| --- | --- | --- | --- | --- | --- | --- | --- | --- |
|  |  |  |  | Mean | Max | Max/mean | Std.dev |  |
| 28 | 10 | signature | 437,863 | 16,900.13 | 1,308,319 | 77.41 | 39,097.01 | 8.65 |
|  |  | universal | 209,314 | 35,353.31 | 2,332,470 | 65.98 | 54,705.09 | 9.30 |
|  |  | frequency | 986,239 | 7503.19 | 197,055 | 26.26 | 8756.40 | 5.50 |
|  |  | universal freq | 214,232 | 34,541.72 | 213,781 | 6.19 | 16,541.84 | 8.47 |
|  | 9 | signature | 123,195 | 60,066.90 | 3,331,373 | 55.46 | 145,924.74 | 9.03 |
|  |  | universal | 51,176 | 144,597.90 | 5,281,396 | 36.52 | 213,163.99 | 9.82 |
|  |  | frequency | 252,085 | 29,354.95 | 212,789 | 7.25 | 33,970.50 | 5.76 |
|  |  | universal freq | 51,591 | 143,434.75 | 331,598 | 2.31 | 67,898.56 | 9.00 |
| 55 | 10 | signature | 212,083 | 22,160.92 | 1,789,219 | 80.74 | 52,883.84 | 16.64 |
|  |  | universal | 120,804 | 38,905.63 | 3,083,513 | 79.26 | 69,992.79 | 16.20 |
|  |  | frequency | 893,645 | 5259.31 | 268,775 | 51.10 | 8764.22 | 11.13 |
|  |  | universal freq | 128,105 | 36,688.31 | 306,121 | 8.34 | 23,739.79 | 15.86 |
|  | 9 | signature | 65,232 | 72,049.84 | 4,827,246 | 67.00 | 187,705.48 | 16.83 |
|  |  | universal | 32,940 | 142,682.31 | 6,766,757 | 47.43 | 258,611.48 | 16.31 |
|  |  | frequency | 231,913 | 20,266.03 | 316,268 | 15.61 | 33,554.57 | 11.34 |
|  |  | universal freq | 34,453 | 136,416.43 | 376,464 | 2.76 | 96,592.41 | 16.07 |

As for the sampling fraction for the frequency counted orderings, in practice we have found that 1% of reads produces fairly consistent results, and can be sampled quickly. Increasing the fraction to 10% had only very minor effects in terms of the metrics we study here.

The frequency ordering reduces the maximum bin size but introduces a large number of very small, fragmented bins (Figure 2). The universal frequency ordering keeps a small maximum bin size, but shifts the low end of the distribution, removing most of the very small bins.

**Figure 2.**
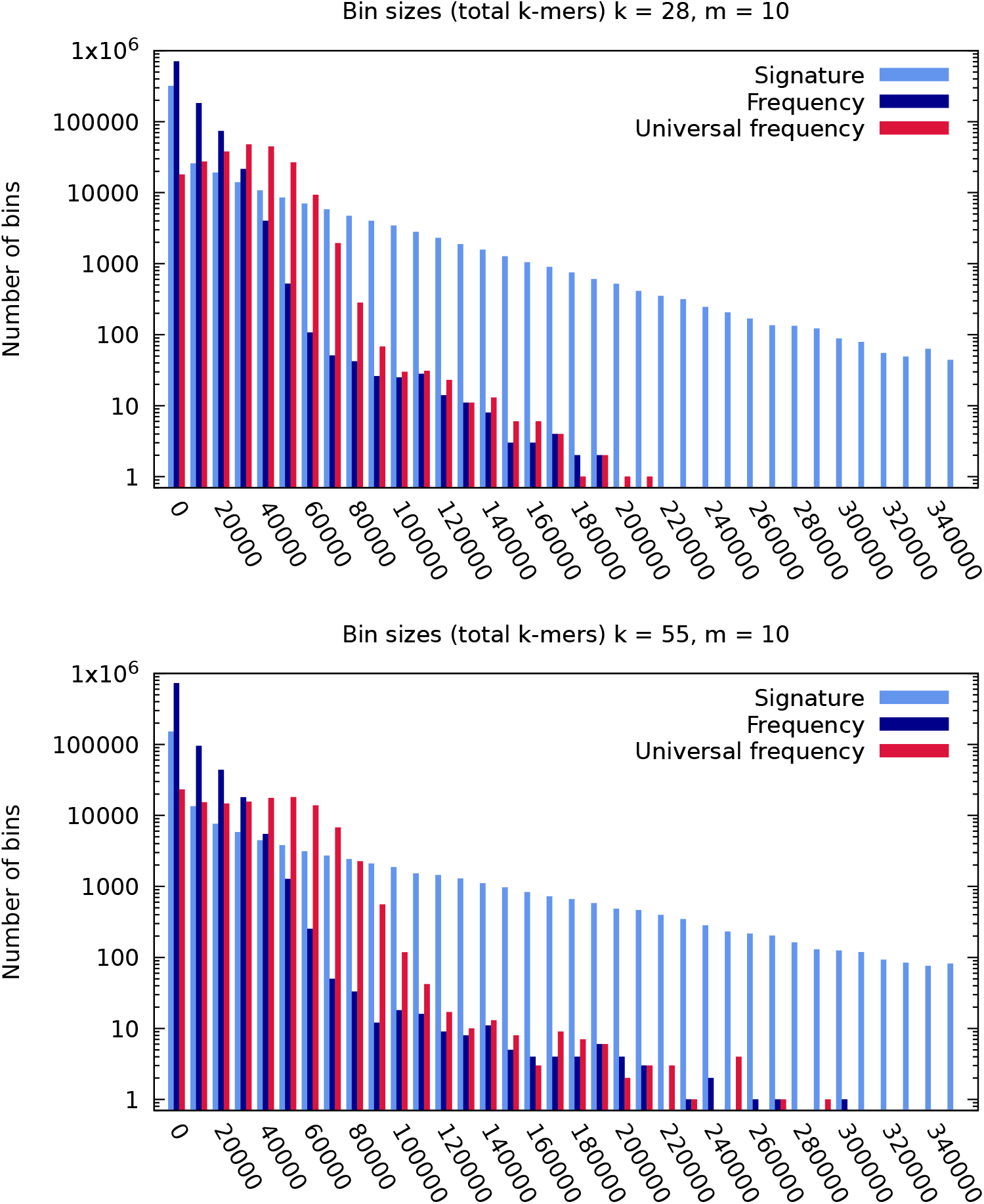
Plot of *k*-mer bin distributions for representative orderings. Note that the x-axis has been cut on the right-hand side. The maxima for all counted and universal counted are present in the charts, but the signature maximum bin is much larger than what this axis can show (Please see Table 3).

### Counting *k*-mers with the universal frequency ordering

Next, we compare the performance of *k*-mer counting on Discount with FastKmer, which was recently introduced by Petrillo et al [8]. FastKmer is a highly efficient distributed *k*-mer counting tool based on Spark, which uses a variant of the signature ordering.

The dataset being studied here is the full data from run SRR094926, cow rumen metagenomics data (see above), from which we previously used only 100 million reads. The size of this dataset is about 314 GB as uncompressed FASTQ files (Table 4).

**Table 4.** Size of the full dataset used for *k*-mer counting.

| $k$ | Total $k$-mers | Distinct $k$-mers |
| --- | --- | --- |
| 28 | $7.23 \times 10^{10}$ | $2.94 \times 10^{10}$ |
| 55 | $4.59 \times 10^{10}$ | $2.50 \times 10^{10}$ |

Benchmarks were run on the Google Cloud Platform (GCP) on three different configurations (Table 5). In each case, four worker machines with sixteen cores each were used. The version of Apache Spark used was 2.4.6. For FastKmer, we used four n1-highmem-16 machines, for a total of 64 CPUs and 416 GB RAM (since FastKmer would not run with less memory). The FastKmer authors’ recommended best settings from [8] were used: *x* = 3*, b* = 8192. However, we increased parallelism (partitions) from the recommended 320 to 2000, since this gave better performance in our setting.

**Table 5.**
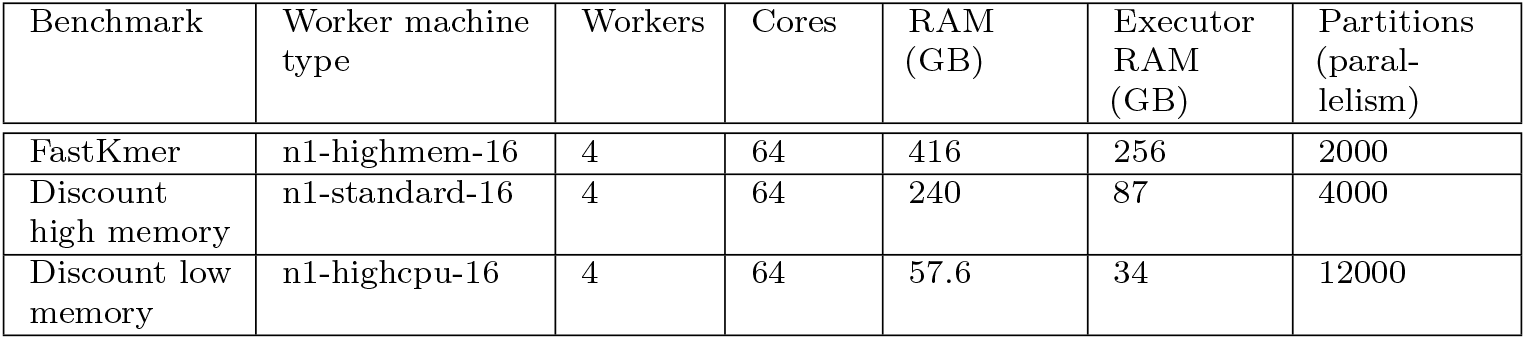
Resource configurations used for performance measurements. Cores and memory (RAM) are reported as totals for all worker machines.

For FastKmer, the number of bins was limited to 8192 as recommended. We also tried larger numbers but did not see a performance improvement. For Discount, the universal frequency ordering was used with the same DOCKS sets as in the previous section, *m* = 10. The number of bins was not constrained and most likely exceeded the number shown in Table 3 (but no greater than the size of the DOCKS sets).

The two applications were performing very similar tasks: outputting a final table with counts for each *k*-mer. Generating this table involves allocating a large amount of strings and writing the data to disk. Hence, this benchmark shows the overall performance effects of nontrivial processing of *k*-mer bins. The outputs from the two applications are identical, except that FastKmer unifies *k*-mers with their reverse complements, while Discount does not.

All machines were from the Google Cloud N1 series, with Intel Xeon CPUs running at 2.7 - 3.2 GHz (all-core turbo frequency). In each case, the cluster master machine was an n1-standard-4 machine with four CPUs.

In a Spark cluster, not all the memory available on the machines is assigned to Spark executors (which run the actual tasks), since some memory is reserved for the operating system, task management, and other functions. For the Discount high memory case, we limited executor memory to test the efficiency of our method. Thus, the total executor memory in that case was only 87 GB across all four machines. For the Discount low memory case, we increased the number of spark partitions to further reduce memory pressure, making them smaller. We also reduced the maximum MapReduce split size (for the underlying file inputs) from the default 128 MB to 16 MB.

We measured the time required by running Discount as well as FastKmer on the full dataset (Table 6). Since FastKmer is internally divided into two main stages, we break down its runtime in the same way as we do for Discount. However, the precise algorithms used by these stages are different between the two applications.

**Table 6.** Performance measurement results. FastKmer does not have a sampling stage, so no timing is reported.

|  |  |  | Runtime (min) |  |  |  |  |
| --- | --- | --- | --- | --- | --- | --- | --- |
| Case | k | m | Sample | Hash | Process | Total | Shuffle (GB) |
| FastKmer | 28 | 10 |  | 21 | 57 | 78 | 125.4 |
|  | 55 | 10 |  | 18 | 43 | 60 | 73.1 |
| Discount<br>high memory | 28 | 10 | 1.3 | 12 | 41 | 55 | 185.7 |
|  | 55 | 10 | 1.3 | 7.8 | 32 | 46 | 96.4 |
| Discount<br>low memory | 28 | 10 | 1.6 | 17 | 55 | 75 | 189 |
|  | 55 | 10 | 1.6 | 10 | 47 | 60 | 100 |

To test scaling to a larger number of worker machines, we also ran the Discount high memory case on sixteen worker machines with four CPUs each. Performance did not change significantly from the four machine case.

## Discussion

In this work, first, we studied various minimizer orderings with the goal of yielding smaller and more evenly distributed *k*-mer binning for a relatively large number of bins.

Four orderings were compared: signature, frequency sampled, universal, and universal frequency sampled. The signature ordering has been a pragmatic choice for many tools since it yields relatively long superkmers and avoids certain large bins.

However, the max/mean ratio remains high, especially for large numbers of bins: up to 80 for *k* = 55, *m* = 10. Figure 2 shows that the bin size distribution for signature is very wide, although nonetheless, smaller bins are more common than large ones.

The universal ordering (lexicographic, DOCKS based) yields even longer superkmers. However, the max/mean ratio improves only slightly relative to the signature case for the same value of *m*. Visually this distribution is very similar to signature, and for simplicity we have chosen not to include it in our charts.

The frequency sampled ordering dramatically reduces the max/mean ratio as well as the size of the maximum bin, but at the expense of a much larger number of bins. As Figure 2 shows, this ordering yields a large amount of very small bins. For some applications, having to maintain so many very small bins will lead to undesirable overhead. Moreover, superkmers are very short, meaning that the bins cannot be stored efficiently in this form.

The universal frequency ordering reduces the max/mean ratio even further and also restores the long superkmers. For a bin number comparable to the basic universal ordering, the max/mean ratio has been reduced by roughly 90%. Superkmers are almost as long as for the signature and universal cases. Thus, for a given number of desired bins, this ordering seems to provide the best balance of long superkmers and a low max/mean ratio. Visually, it is also apparent that the very small bins generated by the frequency sampled ordering are not a problem for this ordering.

Next, we applied the universal frequency ordering to distributed *k*-mer counting, to test a practical application. We evaluated a high memory as well as a low memory scenario while comparing against FastKmer, an existing similar *k*-mer counter. For the former, although the amount of executor memory assigned to Discount is only 34% of what was assigned to FastKmer, Discount runs faster. We believe that this reflects various costs of processing larger bins. For example, both FastKmer and Discount have to sort all *k*-mers in each bin as part of counting, and the cost of sorting increases more than linearly as the array to be sorted grows longer. For the low memory scenario, we carefully tuned Spark to test the limits of our approach. Discount was running on a low memory machine type, with a total of 64 CPUs and only 34 GB total executor memory, around 1/8 of the FastKmer memory. Even with this minimal resource allocation, Discount ran at the same speed as FastKmer in the *k* = 55 case and remained faster for *k* = 28.

Distributed *k*-mer counters can be divided into two categories: out-of-core, (which keep some data on disk) and in-core methods (which keep all data in memory).

FastKmer and Discount are in the former category, since Spark relies on the ability to spill data to disk when necessary. Given the comparison between FastKmer and other tools such as KCH, ADAM and BioPig in [8], which are also out-of-core, this would make Discount both the fastest and the most memory efficient distributed *k*-mer counter in this category. On machines with a given amount of memory, the maximum amount of data that Discount can analyse should be much larger than for comparable tools.

Many tools that use *k*-mer binning have traditionally tried to limit the number of bins. When *k*-mers cannot be evenly distributed, this is reasonable, as increasing the number of bins would also create more noisy, very small bins, introducing overhead. However, with the universal frequency ordering, generating a large number of bins should now be an attractive choice for many applications.

Some open problems and challenges remain. For the universal frequency ordering, we depend on a precomputed universal *k*-mer set. While existing sets work well, unfortunately the DOCKS algorithm for computing new sets can be slow to run, and the authors’ implementation depends on a commercial integer linear programming (ILP) solver tool. However, ongoing work in this area may simplify the generation of such sets in the future. For example, DeBlasio et al have developed an approach for generating such sets by using DOCKS as a starting point and extending them [3]. Other algorithms for generating larger universal sets faster are also being developed [6, 24]. It is also possible to compute species- or genome-specific sets, which may yield even more efficiency [17, 18].

In this work we have only studied selected minimizer orderings of interest, and we leave a broader comparison with other binning methods for future work. For example, Efe [5] suggests a method based on sums of integers associated with the nucleotides of a *k*-mer.

Many *k*-mer processing tools unify each *k*-mer with its reverse complement, treating them as the same value. This is made possible in part by restricting minimizers and superkmers in certain ways. Unfortunately, with our minimizer ordering this is not currently easy to do. This has been recognised as an open problem for universal sets by Marçais et al [15]. In general, research in universal *k*-mer sets is currently ongoing, and recent results may further improve the universal frequency ordering.

## Conclusion

In this work we have investigated the formation of binned superkmers from genomic sequences by using minimizers, a common technique in omics data analysis tools. To support the investigation, we implemented a new distributed *k*-mer counting tool, Discount, which also has minimizer ordering analysis functionality. We sought to achieve an even distribution of bin sizes, with a view to improvements such as memory usage reduction, efficient storage on disk, and reduction of overall processing speed. By combining frequency sampled minimizers with universal *k*-mer sets we effectively obtain universal frequency sampled minimizers. To the best of our knowledge, the present work is the first time this combined ordering has been used. Relative to minimizer signatures, the max/mean ratio for bin sizes was reduced by as much as from 77.41 to 6.19 (for *m* = 10, *k* = 28) while still yielding long superkmers. Furthermore, the cost of sampling is small: for the full dataset, only around 5% of the runtime was spent sampling 1% of the reads. Using Discount, compared with the fastest existing out-of-core distributed *k*-mer counting tool, we were able to count *k*-mers in a metagenomic dataset at the same speed or faster using only 14% of the memory. Considering these benefits, we believe that frequency sampled universal minimizers would significantly improve the performance of many tools that use minimizers to construct binned superkmers, and that this should be a preferred strategy for producing evenly sized bins. With this minimizer ordering, Discount expands the practical boundaries of analysis of very large omics datasets.

## Acknowledgments

J.N.P. is grateful to Shandar Ahmad for discussions and for the invitation to visit Jawaharlal Nehru University, New Delhi, in 2018 through the GIAN programme. The authors are grateful to Yuji Kosugi for comments on the draft.

## Footnotes

1 To see this relationship, consider that the total number of *k*-mers represented by *n* superkmers of length *L* (in base pairs) is *n*(*L* (*k* 1)), since each superkmer has to overlap another by (*k* 1) bps. Thus, the larger *L*, the smaller the fraction of pure overlap data (*k* 1) in the superkmer, and the more efficient the storage.

## References

1. P. A. Audano, S. Ravishankar, and F. O. Vannberg. Mapping-free variant calling using haplotype reconstruction from k-mer frequencies. Bioinformatics, 34(10):1659–1665, 2017.

2. R. Chikhi, A. Limasset, S. Jackman, J. T. Simpson, and P. Medvedev. On the representation of de bruijn graphs. In Lecture Notes in Computer Science (including subseries Lecture Notes in Artificial Intelligence and Lecture Notes in Bioinformatics), volume 8394 LNBI, pages 35–55, 2014.

3. D. DeBlasio, F. Gbosibo, C. Kingsford, and G. Marçais. Practical universal k-mer sets for minimizer schemes. ACM-BCB 2019 - Proceedings of the 10th ACM International Conference on Bioinformatics, Computational Biology and Health Informatics, pages 167–176, 2019.

4. S. Deorowicz, M. Kokot, S. Grabowski, and A. Debudaj-Grabysz. KMC 2: Fast and resource-frugal k-mer counting. Bioinformatics, 31(10):1569–1576, 2015.

5. K. Efe. Robust K-mer partitioning for parallel counting. In BIOINFORMATICS 2018 - 9th International Conference on Bioinformatics Models, Methods and Algorithms, Proceedings; Part of 11th International Joint Conference on Biomedical Engineering Systems and Technologies, BIOSTEC 2018, volume 3, pages 146–153, 2018.

6. B. Ekim, B. Berger, and Y. Orenstein. A Randomized Parallel Algorithm for Efficiently Finding Near-Optimal Universal Hitting Sets. In R. Schwartz, editor, Research in Computational Molecular Biology, pages 37–53, Cham, 2020. Springer International Publishing.

7. M. Erbert, S. Rechner, and M. Muller-Hannemann. Gerbil: A fast and memory-efficient k-mer counter with GPU-support. Algorithms for Molecular Biology, 12(1):1–12, 2017.

8. U. Ferraro Petrillo, M. Sorella, G. Cattaneo, R. Giancarlo, and S. E. Rombo. Analyzing big datasets of genomic sequences: Fast and scalable collection of k-mer statistics. BMC Bioinformatics, 20(Suppl 4):1–14, 2019.

9. M. Hess, A. Sczyrba, R. Egan, T. W. Kim, H. Chokhawala, G. Schroth, S. Luo, D. S. Clark, F. Chen, T. Zhang, R. I. Mackie, L. A. Pennacchio, S. G. Tringe, A. Visel, T. Woyke, Z. Wang, and E. M. Rubin. Metagenomic discovery of biomass-degrading genes and genomes from cow rumen. Science, 331(6016):463–467, 2011.

10. C. Jain, A. Rhie, H. Zhang, C. Chu, B. P. Walenz, S. Koren, and A. M. Phillippy. Weighted minimizer sampling improves long read mapping. Bioinformatics (Oxford, England), 36(1):i111–i118, 2020.

11. H. Karau and R. Warren. High performance spark: best practices for scaling and optimizing Apache Spark. O’Reilly, 2017.

12. M. Kokot, M. Dlugosz, and S. Deorowicz. KMC 3: counting and manipulating k-mer statistics. Bioinformatics (Oxford, England), 33(17):2759–2761, 2017.

13. S. Koren, B. P. Walenz, K. Berlin, J. R. Miller, N. H. Bergman, and A. M. Phillippy. Canu: Scalable and accurate long-read assembly via adaptive κ-mer weighting and repeat separation. Genome Research, 2017.

14. S. C. Manekar and S. R. Sathe. A benchmark study of k-mer counting methods for high-throughput sequencing. GigaScience, 7(12):1–13, 2018.

15. G. Marçais, D. Pellow, D. Bork, Y. Orenstein, R. Shamir, and C. Kingsford. Improving the performance of minimizers and winnowing schemes. In Bioinformatics, volume 33, pages i110–i117, 2017.

16. Y. Orenstein et al. DOCKS public web site, Accessed October 11, 2020. http://acgt.cs.tau.ac.il/docks.

17. Y. Orenstein, D. Pellow, G. Marçais, R. Shamir, and C. Kingsford. Compact universal k-mer hitting sets. In Lecture Notes in Computer Science (including subseries Lecture Notes in Artificial Intelligence and Lecture Notes in Bioinformatics), volume 9838 LNCS, pages 257–268, 2016.

18. Y. Orenstein, D. Pellow, G. Marçais, R. Shamir, and C. Kingsford. Designing small universal k-mer hitting sets for improved analysis of high-throughput sequencing. PLoS Computational Biology, 13(10):1–15, 2017.

19. U. F. Petrillo, G. Roscigno, G. Cattaneo, and R. Giancarlo. FASTdoop: A versatile and efficient library for the input of FASTA and FASTQ files for MapReduce Hadoop bioinformatics applications. Bioinformatics, 33(10):1575–1577, 2017.

20. G. Rizk, D. Lavenier, and R. Chikhi. DSK: K-mer counting with very low memory usage. Bioinformatics, 29(5):652–653, 2013.

21. M. Roberts, W. Hayes, B. R. Hunt, S. M. Mount, and J. A. Yorke. Reducing storage requirements for biological sequence comparison. Bioinformatics, 20(18):3363–3369, 2004.

22. The Apache Software Foundation. Apache Spark, Accessed October 11, 2020. http://spark.apache.org.

23. D. E. Wood and S. L. Salzberg. Kraken: Ultrafast metagenomic sequence classification using exact alignments. Genome Biology, 15(3), 2014.

24. H. Zheng, C. Kingsford, and G. Marçais. Improved design and analysis of practical minimizers. Bioinformatics (Oxford, England), 36(1):i119–i127, 2020.

